# A common marking method influences fish-parasite interactions

**DOI:** 10.64898/2026.10.27.762165

**Authors:** Jessica F. Stephenson

**Affiliations:** Department of Zoology, Stockholm University, Stockholm, Sweden; Department of Biological Sciences, University of Pittsburgh, Pittsburgh, Pennsylvania, USA

## Abstract

Being able to identify the same individual organism through time is essential to robustly address many outstanding questions in ecology and evolution. For organisms that can move, and that have no reliable identifying features, researchers must often use some form of artificial marking system. In fish, a commonly used system is Visible Implant Elastomer (VIE) marking. Previous studies indicate that this marking system does not alter the survival, growth, or behaviour of marked fish. Here, using 26 VIE-marked and 46 control guppies, *Poecilia reticulata*, I found that VIE marking indeed did not impact fish growth. However, one month after marking, VIE-marked fish experienced an estimated ∼3.7-fold higher *Gyrodactylus turnbulli* parasite burden during experimental infections. Two months post-marking, *G. turnbulli* individuals transmited faster to VIE-marked than unmarked fish, although at this timepoint their subsequent growth was not different on fish from different treatment groups. Despite its lack of impact on common metrics of fish health, therefore, VIE marking appears to have important, long-lasting impacts on fish-parasite interactions. These findings suggest caution with assuming that marking animals is biologically neutral: that assumption may fail when the response variable involves host-parasite interactions.

## Introduction

Addressing many outstanding questions in ecology and evolution relies on researchers being able to identify the same individual organism through time. Long-term studies of individually marked animals have been especially powerful because they allow researchers to connect phenotypic variation to survival, reproduction, social interactions, and evolutionary change across time. For example, individually marked Darwin’s finches provided foundational evidence for natural selection in the wild (P. R. Grant et al., 1976), while long-term studies of individually marked Soay sheep have enabled researchers to quantify selection and its relationship with environmental variation across generations (Johnston et al., 2013). More generally, individual identification underpins mark–recapture approaches used to estimate demographic rates, movement, and fitness in natural populations (Lebreton et al., 1992). Insights such as these motivate researchers to overcome the often substantial challenges associated with marking individual animals.

Marking small animals that are mucous-covered or moult frequently presents a special challenge. Among researchers working on such animals, visible implant elastomer (VIE; Northwest Marine Technology, Inc., Anacortes, USA) is a popular choice of mark. Typically, a small bead of inert, coloured polymer is injected under the skin or cuticle. VIE has been evaluated as a mark across animal taxa, including in blow flies, earthworms, termites, reptiles, amphibians, and fish, with results suggesting no impact on survival or development (Bainbridge et al., 2015; Bushon et al., 2007; Butt & Lowe, 2007; Moffatt, 2013; Padget & Thompson, 2021). This last is crucially important for many questions: marking systems must minimally impact the animals so that the data collected accurately represents processes occurring in unmanipulated animals (Batsleer et al., 2020).

Given its popularity as a mark in fish in particular, there is much previous research evaluating the effects of VIE on physiological, morphological, and behavioural measurements of fish across species. Many studies find no significant difference between marked and unmarked fish in measurements such as growth, survival, and social behaviour (Jungwirth et al., 2019; Long et al., 2024; Uglem et al., 2020), supporting the general suitability of VIE for individual identification.

However, some studies have found evidence for potentially important context-specific effects. For example, Jungwirth et al. (2019) found that particular colours of mark could influence social behaviour in cichlids. In addition, Henrich et al. (2014) found that VIE tagging altered the immune response of three-spined sticklebacks.

These findings raise the possibility that VIE could influence biological processes that are not routinely included when validating marking methods, even when commonly measured traits such as survival or growth are unaffected.

Here, I used Trinidadian guppies (*Poecilia reticulata*) to evaluate how VIE marking may affect its interactions with its common parasite, *Gyrodactylus turnbulli*. Guppies are small fish native to freshwater streams in Trinidad, Tobago, and northern South America (Magurran, 2005). VIE is widely used to individually identify guppies in mark–recapture studies, including in long-term experimental populations, which have generated major insights into guppy evolution, demography, and host– parasite interactions (Gordon et al., 2009; Potter et al., 2026; Rogowski et al., 2020; Travis et al., 2023). VIE marks have also been used to follow guppies in natural populations and quantify social relationships and cooperative interactions (Croft, Arrowsmith, et al., 2003; Croft et al., 2006), individual growth and responses to environmental conditions across life stages (Auer et al., 2012), dispersal and gene flow (Fitzpatrick et al., 2016), and the relationship between individual cooperative phenotypes and social assortment (Brask et al., 2019). VIE marking has therefore been instrumental in generating some of the detailed individual-level data that have contributed to the Trinidadian guppy’s status as a model system for studying ecology and evolution.

The extensive use of VIE marking in this system has yielded many tests of its effects on guppy mortality, but fewer tests on other aspects of their biology. Studies often report no mortality (E. Grant et al., 2025; Hansen et al., 2021; Morrell et al., 2008), or up to around 5-7% mortality (Croft, Albanese, et al., 2003; McKellar & Hendry, 2011; Piyapong et al., 2010), and usually also report that fish quickly resume ‘normal swimming behaviour’, although this is not typically quantified. How VIE marking impacts guppy growth has received fewer direct tests: most studies cite Reznick et al. (1996), who report no effects of the marks on mortality or growth over a 12 month period. Similarly, only one study to my knowledge has directly evaluated the effects of VIE on guppy behaviour, and indicates that guppies show no preference for VIE-marked or unmarked conspecifics (Croft et al., 2004). How this marking system may impact guppies beyond their mortality remains uncertain, therefore, with potentially important implications for the hundreds of studies using this mark in this species.

My focus is on one such potential impact of VIE marking: its effect on host– parasite interactions. The guppy and its *Gyrodactylus* spp. parasites are an important model for research in host-parasite interactions (Tepox-Vivar et al., 2022). Understanding the impacts of VIE marking on this interaction is important given the often high prevalence of these parasites in natural guppy populations (Clark et al., 2023; Rovenolt et al., 2024; Stephenson et al., 2015), and the extensive use of VIE marking in field research. *Gyrodactylus* spp. parasites of guppies are directly transmitted viviparous monogenean parasites that live on host skin, have a generation time of 24–36 hours, and can complete their entire lifespan (average 5.5 days) on a single host (Harris & Lyles, 1992; Scott & Nokes, 1984). As such, gyrodactylids can establish substantial infections from just a single worm (Bakke et al., 2007). Because parasite burden and transmission depend on host physiology and immune function, effects of a marking procedure that are undetectable using common measures such as growth or survival could nevertheless alter host–parasite interactions. Here I tested how VIE marks affect the growth, *Gyrodactylus turnbulli* transmission to, and population growth on, 26 marked and 46 unmarked guppies.

## Methods

### Fish origin and maintenance

I used laboratory-bred descendants of wild guppies collected in 2012 from the Caura River in Trinidad (UTM: 20 P 679527.7 m E, 1180376.4 m N based on WGS84 Datum; elevation 112m). All fish were kept on a 12:12 light-dark cycle and fed daily with Tetramin tropical flake and *Artemia* sp. nauplii hatched from decapsulated eggs. Fish were kept on a recirculating system (Aquaneering) with approximately 20% daily water changes (reverse osmosis water, reconstituted with Instant Ocean, magnesium, and calcium) and an average water temperature of 25±1°C.

### Parasite origin and maintenance

The isogenic line of the ectoparasite *Gyrodactylus turnbulli* (species identity confirmed with molecular methods described in (Walsman et al., 2022)) used in this experiment originated from a single parasite isolated from a commercially obtained guppy in 2019. The line was maintained on groups of parasite-naïve fish under standard conditions (hereafter, ‘culture fish’). The number of parasites on each culture fish was counted biweekly (*G. turnbulli* infects guppy skin), and more naïve fish were added as necessary to maintain parasite numbers.

### Visible Implant Elastomer (VIE) injection

I injected 26 guppies (13 females and 13 males) with white visible implant elastomer (hereafter VIE, Northwestern Marine supplies) following protocols similar to those employed by others using these marks in Trinidadian guppies (Fitzpatrick et al., 2013; Travis et al., 2023). I anesthetized guppies with MS-222 (ethyl 3– aminobenzoic methanesulfonate salt, buffered to neutral pH with sodium bicarbonate) and injected the elastomer marks next to the dorsal fin. I handled 46 control fish (21 females and 25 males) similarly with regards to the anaesthetic, but did not inject them.

Marking was conducted at two timepoints: data from these two timepoints constitutes my ‘lag’ treatment group comparison. The first group of fish (7 marked, 9 control fish) was marked in January of 2020, and the second (19 marked, 37 control fish) in December 2020. In each case, once they had been marked, I held the guppies in 1.8L tanks individually before infecting them. The first group were held for one month between injection and infection, and the second group for two months.

While I maintained as many things as possible consistent across these groups, some factors were beyond my control and may have impacted how the host and parasite interacted. For example, there may have been evolution in the parasite, or slight changes in water chemistry or microbiome, which can change the rate at which gyrodactylids grow on their hosts (Gheorghiu et al., 2007, 2012; Kramp et al., 2026; Walsman et al., 2022). Importantly, however, marked and unmarked fish were included at each timepoint, and the comparison between them is the focus of my analysis.

### Infection and monitoring

I manually infected the 72 guppies (34 females, 38 males) following established protocols in this system. I placed anesthetized (buffered MS-222, as above) guppies next to an anesthetized “donor” fish until two parasites had moved from the donor to the experimental recipient, as observed under a dissecting scope. For a subset of the fish in lag group 2 (n=48), I recorded the number of seconds it took for the two parasites to move from donor to recipient. After infection, I returned the experimental fish to their individual 1.8 L tanks under the standard conditions described above.

I quantified how well the parasite populations grew on fish in the different treatment groups. Every other day, I anaesthetised each fish (buffered MS-222) and counted the number of parasites on the body and fins under a dissection microscope. I continued counting the worms on the fish for the duration of the fish’s infection, up to a maximum of 22 days.

### Length and weight data

I collected weight and length data at three timepoints in the experiment: before injection, before infection, and after infection. Fish were anaesthetised (buffered MS-222) and placed gently on a folded, moistened, paper towel. I measured their standard length to the nearest 0.1mm using callipers and then weighed them to the nearest 0.001g. I then returned the fish to their individual holding tanks. The fish in lag group 1 were only measured before and after infection, with some fish missing these data. The fish in lag group 2 were measured at all three timepoints, unless they died before the ‘after infection’ timepoint.

### Data analysis

I used the data to address three questions: 1) Does marking treatment affect fish growth?; 2) Do parasites transmit differently to fish in different marking treatment groups?; 3) Do parasites grow differently on fish in different marking treatment groups? All analyses to address these questions were conducted in R version 4.4.0 (R Core Team, 2023), and the data, code, and output are in the supplementary material. I validated the fit of each of my statistical models (described below) using the *DHARMa* package (Hartig, 2022), and provide the plots in the supplementary material. I used *emmeans* (Lenth, 2024) to extract effect sizes, and *ggplot2* (Wickham, 2016) for the figures.

To address the first question, I tested how the treatment groups differed in their change in length and weight across each of the two time periods: post injection (from pre-injection to pre-infection timepoints); and post infection (from pre-infection to post-infection timepoints). I used four linear models, each of which used weight/length at the later time point as the response variable, with weight/length at the earlier time point, treatment (VIE *vs*. CONTROL), and sex as fixed effects. These models revealed no significant difference between treatment groups. To evaluate the reliability of this conclusion, I conducted a power analysis.

To address the second question, I used parasite transmission latency in seconds as the response variable in a linear mixed effects model using the *lme4* package (Bates et al., 2015). This model included treatment (VIE *vs.* CONTROL), host sex, and donor sex as fixed effects, and donor identity as a random effect as each donor (n=13) was used to infect multiple experimental fish. Because each donor infected experimental fish haphazardly assigned to each of the treatment groups, I conducted a ‘leave one out’ analysis to evaluate how sensitive my results were to the presence of each donor in the dataset.

To address the third question, I first used the number of worms on each fish on each day of its infection to calculate its overall ‘infection severity’ as the area under the curve. I used this infection severity metric as the response variable in a generalised linear model (GLM) with a Gamma error distribution and log link function. As fixed effects, I included treatment (VIE *vs.* CONTROL), lag (one *vs.* two months between injection and infection), sex, the duration of that fish’s infection in days, and the interaction between treatment and lag. Because of my modest sample size, particularly in lag group 1, I conducted a ‘leave one out’ analysis to evaluate how sensitive my results were to the presence of each individual fish in the dataset.

## Results

There was no difference between VIE-marked and control fish in the change in either length or weight across either of the two time periods (Fig. 1, all *P* > 0.09; see supplement). Unsurprisingly, given that female guppies have indeterminate growth and males determinate, sex was a significant predictor of growth in three of these four models (see supplementary material for details). The power analysis showed that I had over 80% power to detect 5% difference in length between treatment groups, and 11% difference in weight (Fig. S2), likely due to a smaller sample size for weight measurements. Because 11% is a substantial difference in weight, and smaller differences are ecologically important (Abbott et al., 1985; Alward et al., 2021; Gilmour et al., 2005), I focus on the analysis of length here (but present equivalent results for weight in the supplement).

**Figure 1.**
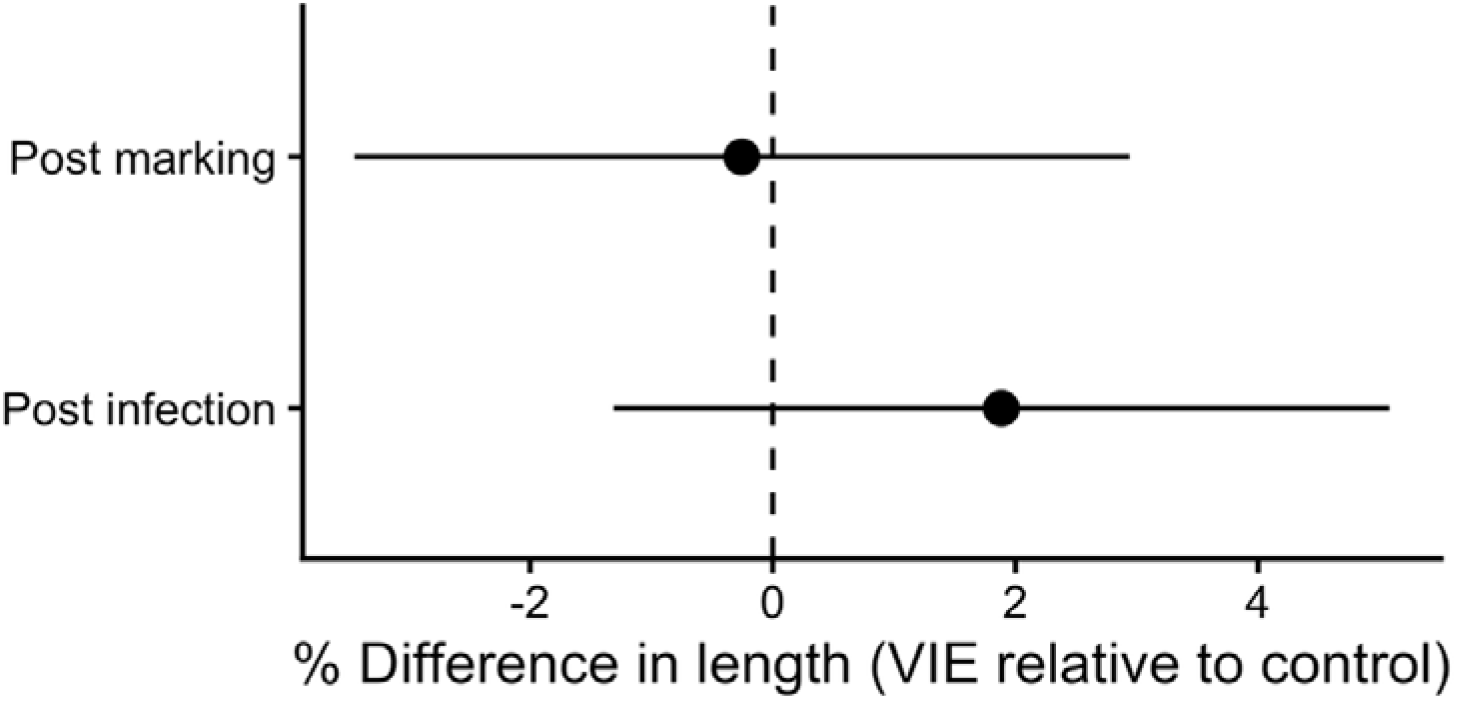
VIE marking does not affect guppy length change over the experimental periods. The points gives the model estimated marginal means converted to percentage change in length between the pre-injection and pre-infection timepoints (‘post marking’), and between the pre- and post-infection timepoints (‘post infection’). The lines gives the 95% confidence intervals.

*Gyrodactylus turnbulli* tended to transmit with lower latency to fish that had been VIE-marked (Fig. 2; χ*²* = 4.46, df=1, *P* = 0.035). No other fixed effect explained variation in transmission latency. Note that all of the data for this test came from fish in lag group 2. Donor identity explained ∼24% of the variance in parasite transmission latency. In line with this finding, a leave-one-donor-out sensitivity analysis showed that the estimated latency was consistently shorter in VIE-marked fish, with estimated reductions ranging from 24% to 56% across the 13 analyses.

**Figure 2.**
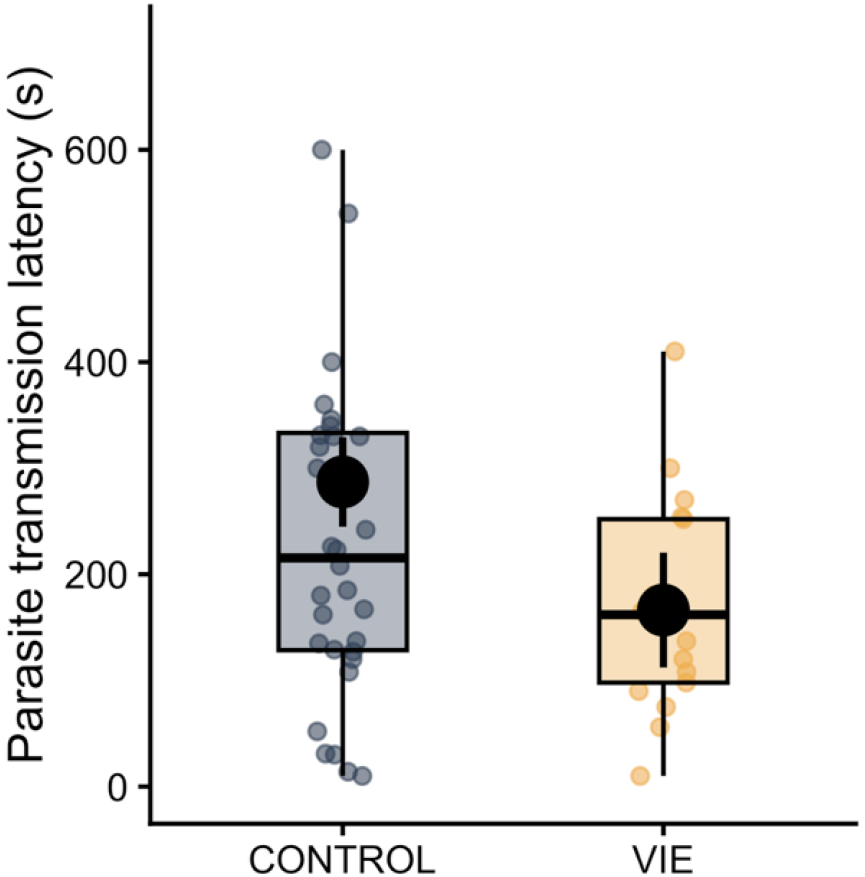
*Gyrodactylus turnbulli* transmit significantly faster to VIE-marked than control fish, 2 months after injection. The coloured points give the raw data of the parasite’s latency to transmit to each experimental fish. The thick line gives the median value, the boxes the interquartile range, and the whiskers the data within 1.5x of the interquartile range. The black points and error bars give the model-estimated marginal means and standard errors.

However, removing individual donors resulted in the treatment effect no longer being statistically significant in 5 of the 13 analyses.

I found that the effect of VIE marking on parasite growth differed between fish that had a lag of 1 month between injection and infection and those that had a lag of 2 months (Fig. 3; lag:treatment interaction: χ*²* = 6.28, df=1, *P* = 0.012). The model estimated that one month after marking, VIE-marked fish had a 3.7-fold higher parasite burden across the infection period than unmarked controls (Fig. 3; SE=1.78, df=63, t=2.71, *P* = 0.009). In contrast, no evidence of a difference was detected two months after marking (VIE:CONTROL ratio = 0.81, SE=0.22, df=63, t=-0.78, *P* = 0.44). The leave one out showed that these results are not sensitive to the presence of any one fish. At one month post-marking the VIE-to-control infection severity ratios ranged from 2.63 to 4.92, and the lower confidence limit was always above 1: VIE marking was associated with higher infection intensities across leave-one-fish-out analyses. By contrast, at two months post-marking, ratios ranged from 0.68 to 0.88, and the confidence interval always included 1: the effect of VIE marking on infection intensity was not significant in any of the leave-one-fish-out analyses.

**Figure 3.**
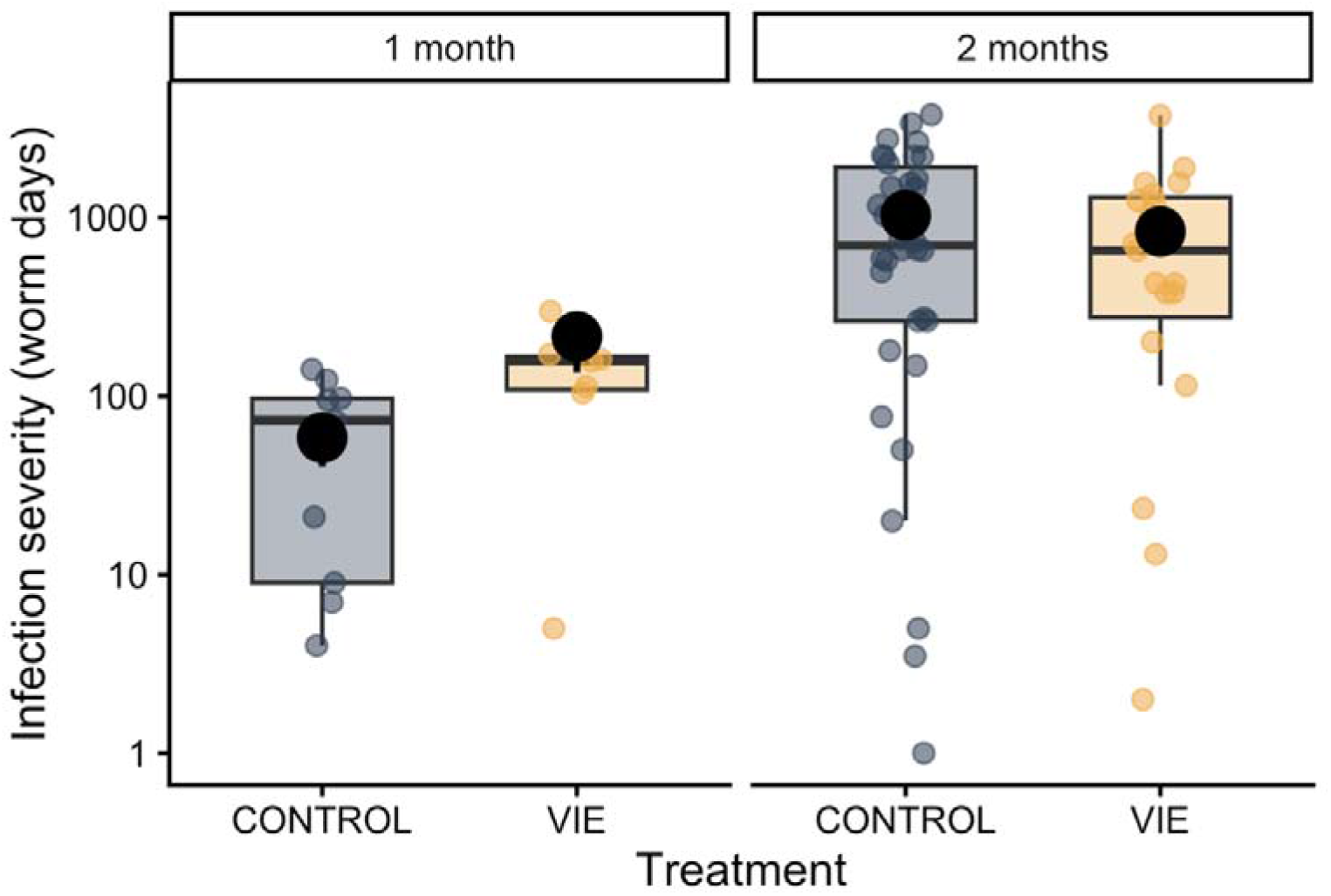
VIE marking significantly impacted *Gyrodactylus turnbulli* growth on guppies 1 month after injection. There was no significant difference in parasite growth on guppies 2 months after injection. The coloured points give the raw data of each fish’s infection severity (quantified as the area under the curve of parasite load over time). The thick line gives the median value, the boxes the interquartile range, and the whiskers the data within 1.5x of the interquartile range. The black points and error bars give the model-estimated marginal means and standard errors.

## Discussion

I found that VIE marking did not affect fish growth, in line with previous research, but it did affect how guppies interacted with their gyrodactylid parasites. One month after marking, VIE-marked fish experienced more severe infections than unmarked controls. Even two months after marking, the parasite transmitted more readily to VIE-marked fish than to unmarked controls, though its subsequent growth did not differ between these groups. These results suggest that, even while not affecting host growth, the effects of VIE marking on host–parasite interactions are long lasting and have potentially important implications for studies using this marking system.

Here I place these results in the context of previous work on this host-parasite system, discuss limitations of the study, and end with the broader implications.

The results suggest that VIE marking can affect both guppy probability of becoming infected, and parasite load once infected, which previous work demonstrates can substantially affect guppy survival, reproduction, and social behaviour. The magnitude of these effects could matter disproportionately in nature. *Gyrodactylus* spp. transmission occurs during contact between infected and susceptible fish, so a change in the time required for parasites to establish on a recipient determine whether or not transmission occurs during a finite period of contact (Tepox-Vivar et al., 2022). As well as binary infection probability, VIE marking resulted in an approximately 9-fold higher parasite burden. In this system, as in many others, the effects of infection on survival (Evanov et al., 2026; van Oosterhout et al., 2007), reproduction (Evanov et al., 2026; Weiler et al., 2025), and behaviour (Jog et al., 2022; Stephenson, 2019; Stephenson et al., 2018) depend on parasite burden. Thus, VIE marking may, through increasing guppy susceptibility to *G. turnbulli* infection, ultimately impact these key metrics in natural settings where these parasites are highly prevalent (Gotanda et al., 2013; Mohammed et al., 2020; Stephenson et al., 2015).

One limitation of this study is that the infections of the two lag groups were conducted several months apart, thus the differences in how well the parasite grew on fish at these timepoints could be due to several factors, beyond the length of time since VIE injection. Importantly, as both VIE-marked and control fish were included at each timepoint, this limitation should not affect the interpretation of the overall effect of VIE marking on this host-parasite interaction. *Gyrodactylus* spp. growth is influenced by a range of host, parasite and environmental factors, including temperature, host size and condition, sex, host immune history and parasite lineage (Harris & Lyles, 1992; Scott & Nokes, 1984; Tepox-Vivar et al., 2022). While I controlled as many factors as I could between these timepoints, the overall difference in parasite growth between these cohorts cannot be attributed to the time since VIE injection. Nevertheless, the fact that the VIE-associated difference in parasite growth was absent after two months, while a difference in transmission latency was still detectable, is consistent with the possibility that effects of marking change through time and may affect different components of the host–parasite interaction for different durations.

The comparison here evaluates the VIE marking procedure as it is routinely conducted in guppy research, in which anaesthetised fish receive an injection of coloured elastomer. The anaesthesia-only control is therefore informative about the net effect of the procedure that field researchers employ, even though it does not isolate the elastomer bead from the needle wound. A sham-injection control (needle insertion with saline) was not included, so the present design cannot distinguish effects attributable to the elastomer material from those attributable to the injection wound itself. Future work should include this arm to decompose the procedure into its mechanistic components. The VIE marking procedure, which combines a needle injection with deposition of an elastomer bead, has important, long-lasting impacts on fish-parasite interactions; whether these effects are driven by the elastomer material, the injection wound, or both remains to be determined.

One possible mechanism for the effect of VIE injection on *G. turnbulli* growth, and apparent latency to colonise a new host, is that VIE injection alters the physical and chemical environment of the skin. Infection itself induces rapid changes in guppy epidermal thickness and mucous-cell characteristics, and experimentally altering the host environment with zinc changes both these epidermal responses and parasite population growth (Gheorghiu et al., 2007, 2012). VIE injection creates a local wound and introduces a foreign material beneath the skin, so it could plausibly alter epidermal structure, mucus production, or other properties of the host surface in ways that influence parasite attachment or reproduction, such as the host skin-associated microbiome (Kramp et al., 2026). A second possibility is altered host immunity. Henrich et al. (2014) found that VIE tagging and associated handling altered immune measures in three-spined sticklebacks, although their sham treatment also produced an immune response and the authors recommended a minimum two-week recovery period. My finding of a parasite effect one month after marking suggests that any transient physiological response to marking could persist longer than is apparent from conventional welfare measures, or could interact with the subsequent infection in a host-specific way. The attenuation of the parasite-growth effect after two months is compatible with such a mechanism.

If VIE marking alters host resistance or parasite establishment, it could introduce systematic variation into studies of host–parasite interactions. Although my results do not establish whether such effects occur in other species or parasite systems, or persist beyond the time periods examined here, they demonstrate that validating a marking method against survival, growth and behaviour does not necessarily establish its biological neutrality for all response variables. The appropriate recovery period after marking may therefore depend on the biological question being asked, particularly in ecological and conservation studies where even small changes in infection or survival could have consequences at individual or population levels (Long et al., 2024). VIE remains a valuable tool for longitudinal studies, but these results should remind researchers to directly evaluate its effects on focal outcomes.

## Acknowledgements

I thank Emmalina Calcaterra, David R. Clark, Lindsay Colgan, Mary J. Janecka, Rachael D. Kramp, Regina Patrick, Maura Sackett, and Jason C. Walsman for technical assistance.

## AI declaration

I used ChatGPT to edit and draft code for my analyses and figures, and Google Scholar Labs to identify relevant literature. All interpretation, writing, scientific content, and conclusions are my own.

## Data accessibility

My data and code are in the supplementary material.

## Ethics

This work was approved by the University of Pittsburgh’s Institutional Animal Care and Use Committee (IACUC), protocol 21069471.

