## supplementary material for "A common marking method influences fish-parasite interactions"

Jessica F. Stephenson

**5 R and Library Versions** **23**

### 1 Data description

Table S1: README of the variables in the dataset

| Variable | Description |
|---|---|
| fishID | fish identity |
| sex | sex of the fish; M=male, F=female |
| treat | treatment group; VIE=fish injected with visible implant elastomer mark, CONTROL=fish not injected |
| lag | the time lag between injection treatment and infection, in months |
| firstw | weight of the fish before injection (mg) |
| firstl | length of the fish before injection (mm) |
| batch | the batch of the infection: 1-4 |
| timeinfs | the number of seconds it took for the parasites to transmit to the experimental host during manual infection |
| prew | weight of the fish before infection (mg) |
| prel | length of the fish before infection (mm) |
| donorid | identity of the donor that was used to infect the experimental fish |
| donorsex | sex of the donor that was used to infect the experimental fish |
| latew | weight of the fish after infection (mg) |

Table S1: README of the variables in the dataset *(continued)*

| Variable | Description |
|---|---|
| latel | length of the fish after infection (mm) |
| day | day of infection |
| dose | the number of worms that established the infection on the experimental fish |
| totworm | the total number of worms on the fish |

##### 1.1 Data management

```
df$fishID <- factor(df$fishID)
df$treat <- factor(df$treat)
df$lag <- factor(df$lag)
df$sex <- factor(df$sex)
df0<-subset(df, day>0)

#Subset down to one row per fish
fdf <- df %>%
  group_by(fishID) %>%
  slice(1) %>%
  ungroup()

fdf2<-subset(fdf, lag==2)
```

#### 2 Testing how marking affects change in length and weight

##### 2.1 Models

Here I run four separate models testing how length and weight change after injection, and after infection. None of them find a significant effect of marking treatment.

```
m_weight_preinfection <- lm(prew ~ treat + firstw + sex,
  data = fdf2
)
Anova(m_weight_preinfection)
```

```
## Anova Table (Type II tests)
##
## Response: prew
##             Sum Sq Df  F value  Pr(>F)
## treat     0.000665  1   2.9904 0.09019 .
## firstw    0.034119  1 153.4117 < 2e-16 ***
## sex       0.001425  1   6.4092 0.01468 *
## Residuals 0.010675 48
## ---
## Signif. codes:  0 '***' 0.001 '**' 0.01 '*' 0.05 '.' 0.1 ' ' 1
```

```
summary(m_weight_preinfection)
```

```
##
## Call:
## lm(formula = prew ~ treat + firstw + sex, data = fdf2)
##
```

```
## Residuals:
##       Min        1Q    Median        3Q       Max
## -0.036250 -0.006523  0.001000  0.007448  0.042250
##
## Coefficients:
##              Estimate Std. Error t value Pr(>|t|)
## (Intercept)  0.034816   0.010896   3.195  0.00247 **
## treatVIE    -0.007508   0.004342  -1.729  0.09019 .
## firstw       0.755950   0.061033  12.386  < 2e-16 ***
## sexM        -0.017323   0.006842  -2.532  0.01468 *
## ---
## Signif. codes:  0 '***' 0.001 '**' 0.01 '*' 0.05 '.' 0.1 ' ' 1
##
## Residual standard error: 0.01491 on 48 degrees of freedom
##   (4 observations deleted due to missingness)
## Multiple R-squared:  0.9228, Adjusted R-squared:  0.918
## F-statistic: 191.3 on 3 and 48 DF,  p-value: < 2.2e-16
```

```
confint(m_weight_preinfection, "treatVIE")
```

```
##                 2.5 %      97.5 %
## treatVIE -0.01623705 0.001221538
```

```
simf<-simulateResiduals(m_weight_preinfection, plot=T)
```

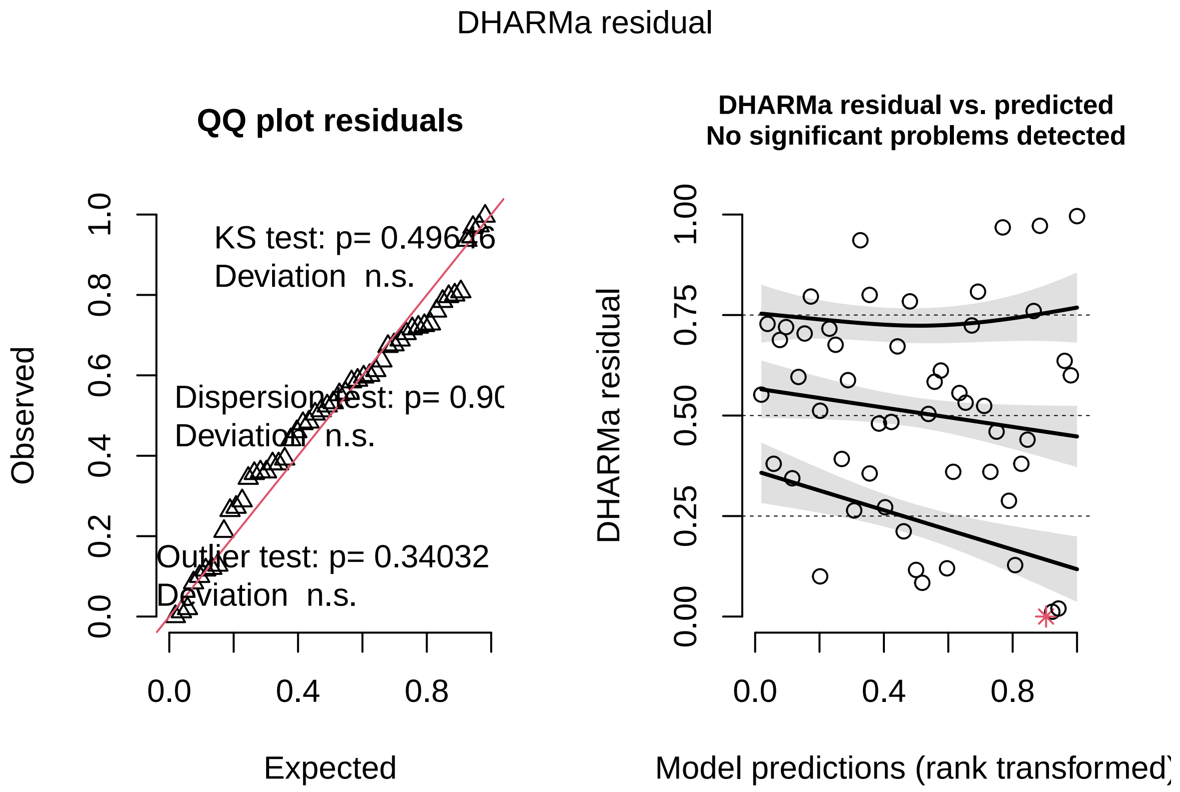

```
testDispersion(simf)
```

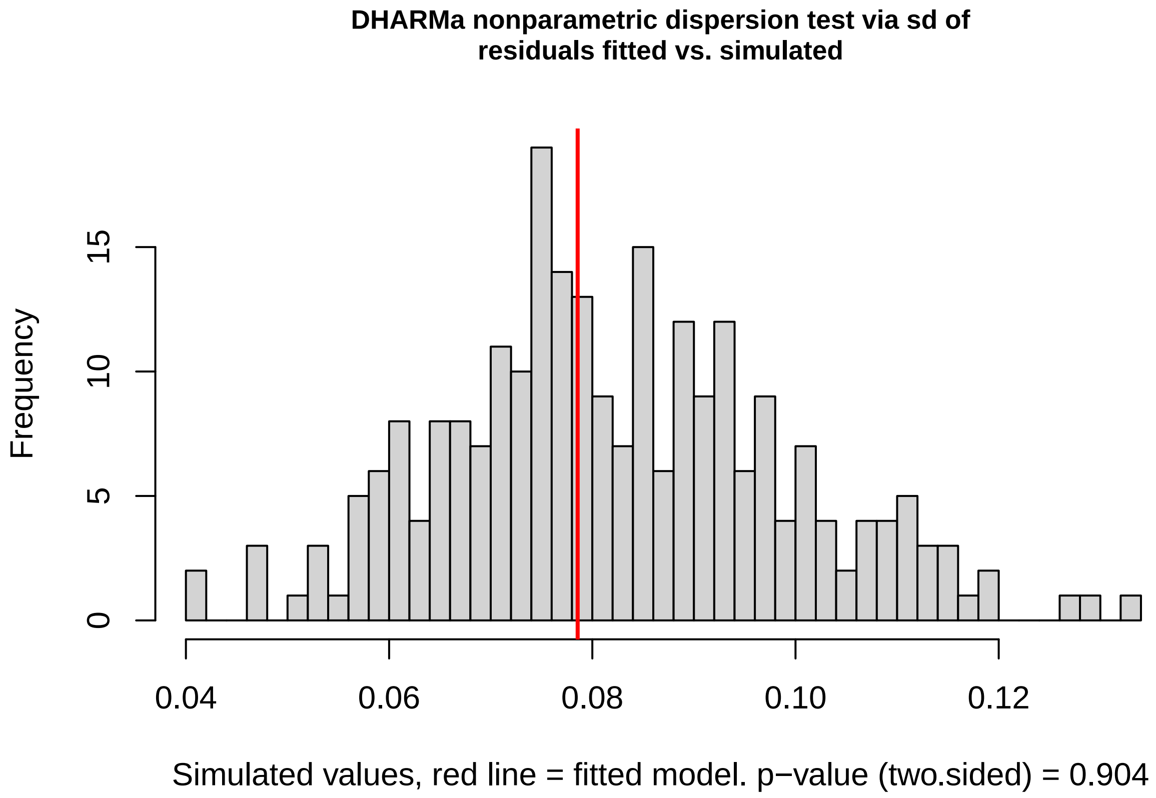

```r
m_weight_infection <- lm(latew ~ treat + prew + sex,
  data = fdf2)
Anova(m_weight_infection)
```

```
## Anova Table (Type II tests)
##
## Response: latew
##             Sum Sq Df  F value    Pr(>F)
## treat     0.000191  1   0.8263    0.3692
## prew      0.037689  1 163.4297 3.821e-15 ***
## sex       0.000018  1   0.0765    0.7837
## Residuals 0.008533 37
## ---
## Signif. codes:  0 '***' 0.001 '**' 0.01 '*' 0.05 '.' 0.1 ' ' 1
```

```r
summary(m_weight_infection)
```

```
##
## Call:
## lm(formula = latew ~ treat + prew + sex, data = fdf2)
##
## Residuals:
##       Min        1Q    Median        3Q       Max
## -0.034906 -0.010286  0.001669  0.007788  0.033714
##
## Coefficients:
##              Estimate Std. Error t value Pr(>|t|)
## (Intercept)  0.006860   0.012498   0.549    0.586
## treatVIE     0.004423   0.004865   0.909    0.369
## prew         0.940810   0.073593  12.784 3.82e-15 ***
```

```
## sexM         -0.002143   0.007750  -0.277    0.784
## ---
## Signif. codes:  0 '***' 0.001 '**' 0.01 '*' 0.05 '.' 0.1 ' ' 1
##
## Residual standard error: 0.01519 on 37 degrees of freedom
##   (15 observations deleted due to missingness)
## Multiple R-squared:  0.9211, Adjusted R-squared:  0.9147
## F-statistic:   144 on 3 and 37 DF,  p-value: < 2.2e-16
```

```r
confint(m_weight_infection, "treatVIE")
```

```
##                  2.5 %     97.5 %
## treatVIE -0.005435474 0.01428091
```

```r
simf<-simulateResiduals(m_weight_infection, plot=T)
```

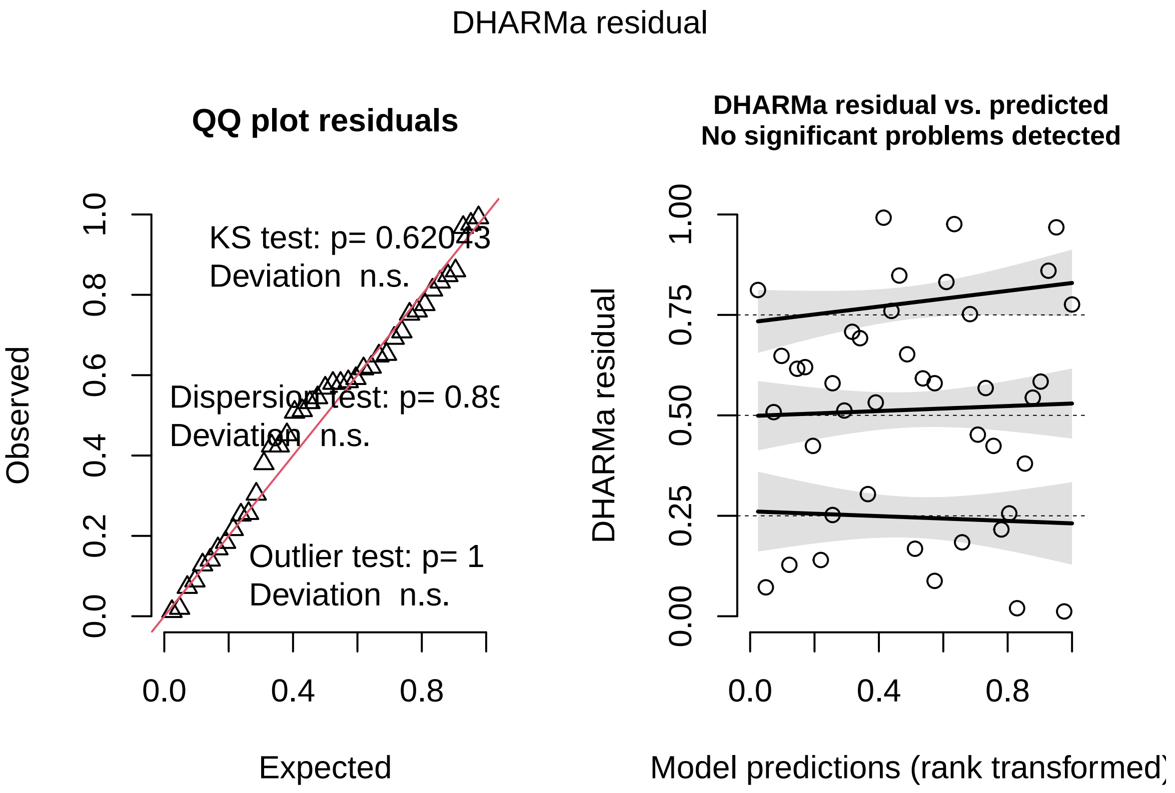

```r
testDispersion(simf)
```

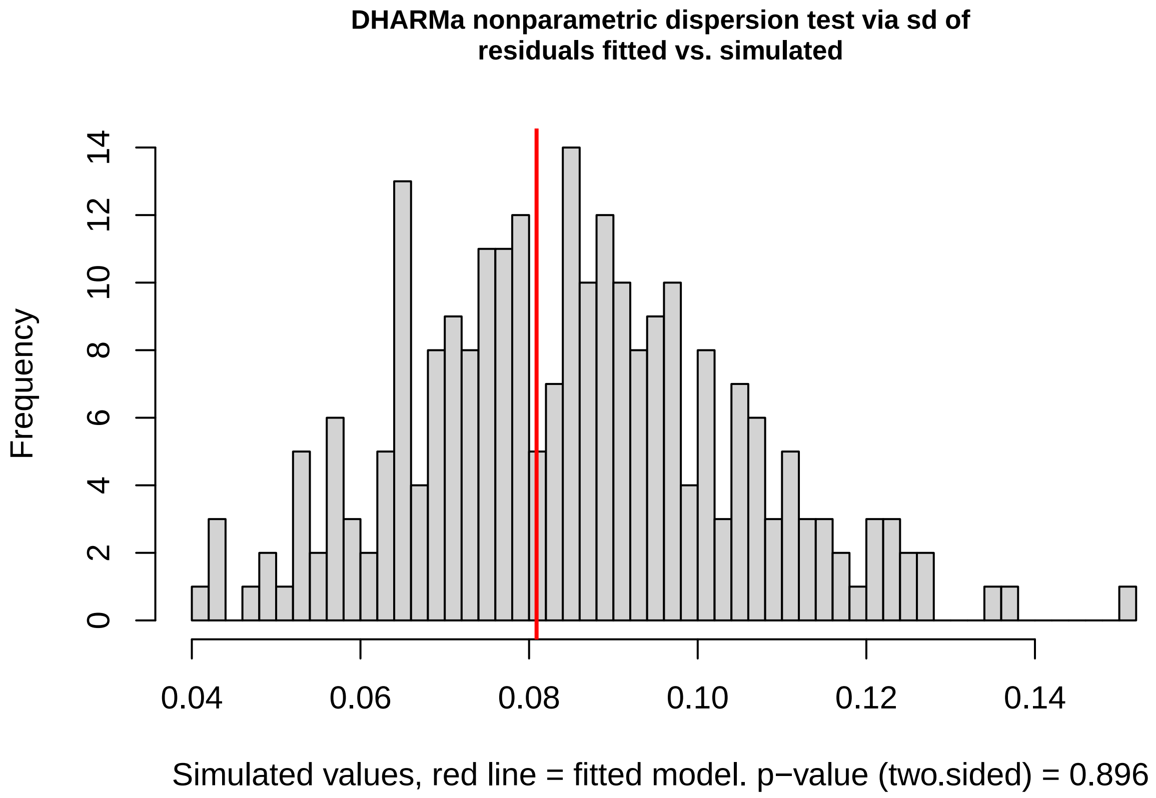

```r
m_length_preinfection <- lm(prel ~ treat + firstl + sex,
  data = fdf2
)
Anova(m_length_preinfection)
```

```
## Anova Table (Type II tests)
##
## Response: prel
##           Sum Sq Df F value    Pr(>F)
## treat      0.025  1  0.0256   0.87354
## firstl    64.395  1 65.3665 1.639e-10 ***
## sex        3.195  1  3.2429   0.07802 .
## Residuals 47.286 48
## ---
## Signif. codes:  0 '***' 0.001 '**' 0.01 '*' 0.05 '.' 0.1 ' ' 1
```

```r
summary(m_length_preinfection)
```

```
##
## Call:
## lm(formula = prel ~ treat + firstl + sex, data = fdf2)
##
## Residuals:
##     Min      1Q  Median      3Q     Max
## -1.8283 -0.6692 -0.1033  0.5869  2.3450
##
## Coefficients:
##             Estimate Std. Error t value Pr(>|t|)
## (Intercept)  4.82548    1.90978   2.527   0.0149 *
## treatVIE    -0.04603    0.28767  -0.160   0.8735
```

```
## firstl        0.78123    0.09663   8.085 1.64e-10 ***
## sexM         -0.90148    0.50060  -1.801   0.0780 .
## ---
## Signif. codes:  0 '***' 0.001 '**' 0.01 '*' 0.05 '.' 0.1 ' ' 1
##
## Residual standard error: 0.9925 on 48 degrees of freedom
##   (4 observations deleted due to missingness)
## Multiple R-squared:  0.8654, Adjusted R-squared:  0.857
## F-statistic: 102.8 on 3 and 48 DF,  p-value: < 2.2e-16
```

```
confint(m_length_preinfection, "treatVIE")
```

```
##                2.5 %    97.5 %
## treatVIE -0.6244321 0.5323698
```

```
simf<-simulateResiduals(m_length_preinfection, plot=T)
```

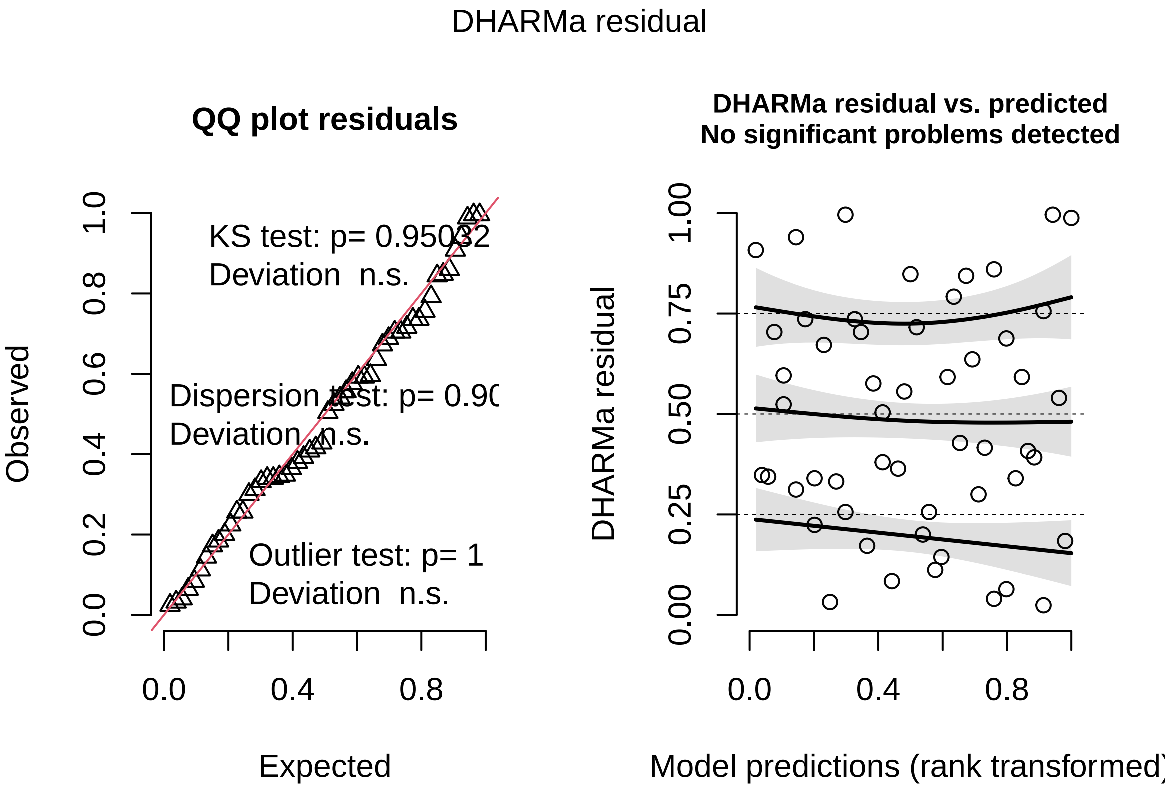

```
testDispersion(simf)
```

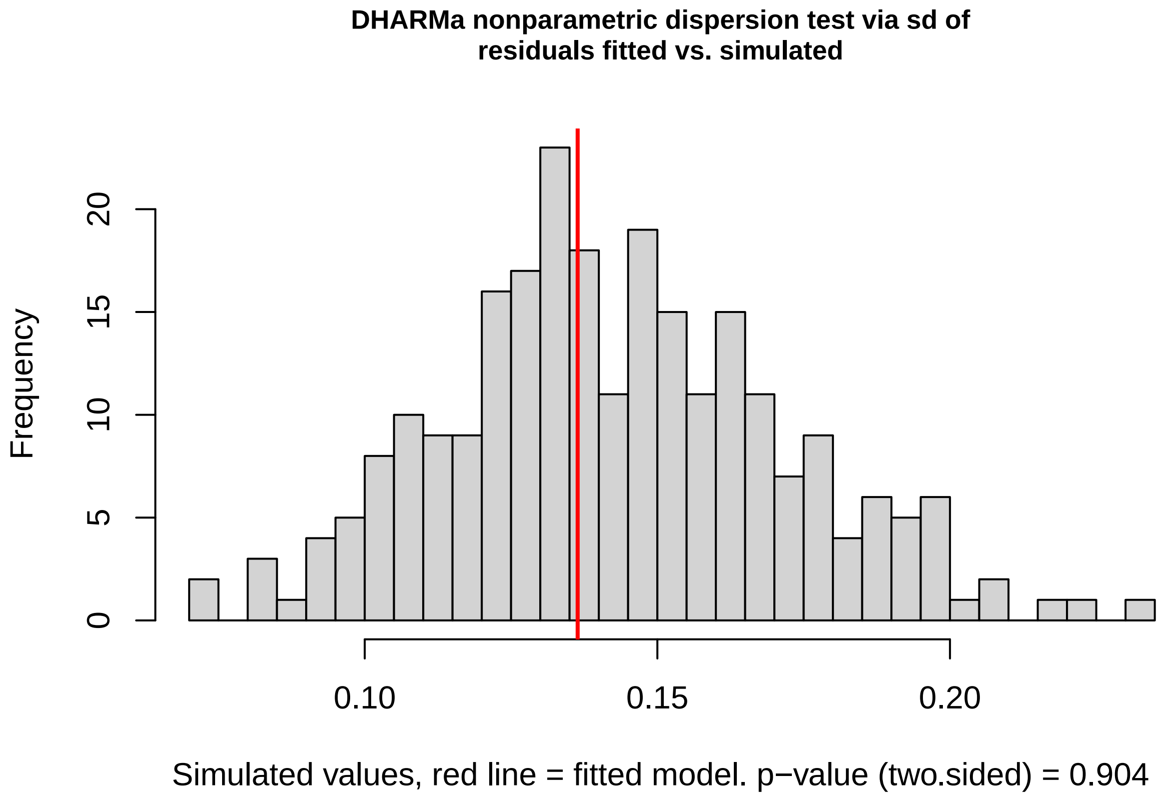

```r
m_length_infection <- lm(latel ~ treat + prel + sex,
  data = fdf2
)
Anova(m_length_infection)
```

```
## Anova Table (Type II tests)
##
## Response: latel
##           Sum Sq Df F value    Pr(>F)
## treat      1.206  1  1.4265   0.23993
## prel      60.190  1 71.2043 3.788e-10 ***
## sex        3.664  1  4.3348   0.04431 *
## Residuals 31.276 37
## ---
## Signif. codes:  0 '***' 0.001 '**' 0.01 '*' 0.05 '.' 0.1 ' ' 1
```

```r
summary(m_length_infection)
```

```
##
## Call:
## lm(formula = latel ~ treat + prel + sex, data = fdf2)
##
## Residuals:
##     Min      1Q  Median      3Q     Max
## -2.6917 -0.4628  0.1207  0.4377  2.0979
##
## Coefficients:
##               Estimate Std. Error t value Pr(>|t|)
## (Intercept)  4.14227    1.95661   2.117   0.0410 *
## treatVIE     0.35541    0.29757   1.194   0.2399
```

```
## prel         0.81869    0.09702   8.438 3.79e-10 ***
## sexM        -1.03153    0.49545  -2.082   0.0443 *
## ---
## Signif. codes:  0 '***' 0.001 '**' 0.01 '*' 0.05 '.' 0.1 ' ' 1
##
## Residual standard error: 0.9194 on 37 degrees of freedom
##   (15 observations deleted due to missingness)
## Multiple R-squared:  0.8908, Adjusted R-squared:  0.8819
## F-statistic: 100.6 on 3 and 37 DF,  p-value: < 2.2e-16
```

```r
confint(m_length_infection, "treatVIE")
```

```
##                2.5 %    97.5 %
## treatVIE -0.2475194 0.9583378
```

```r
simf<-simulateResiduals(m_length_infection, plot=T)
```

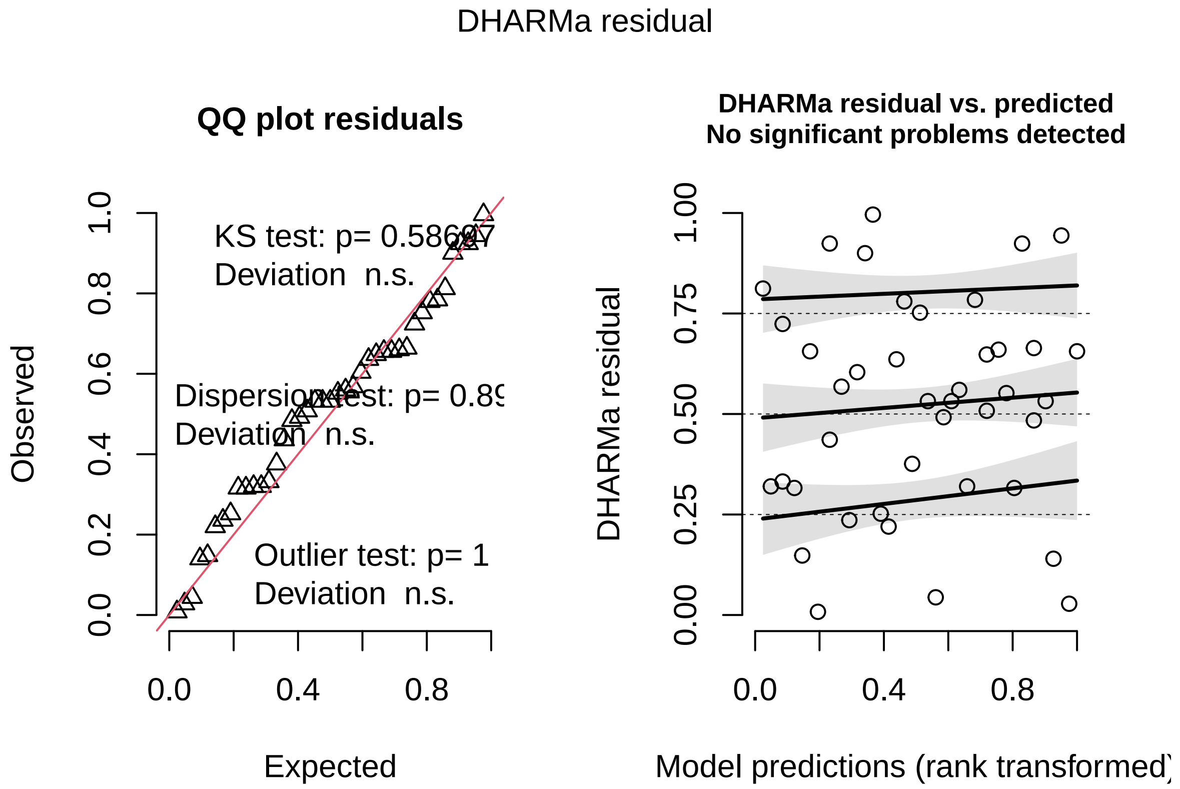

```r
testDispersion(simf)
```

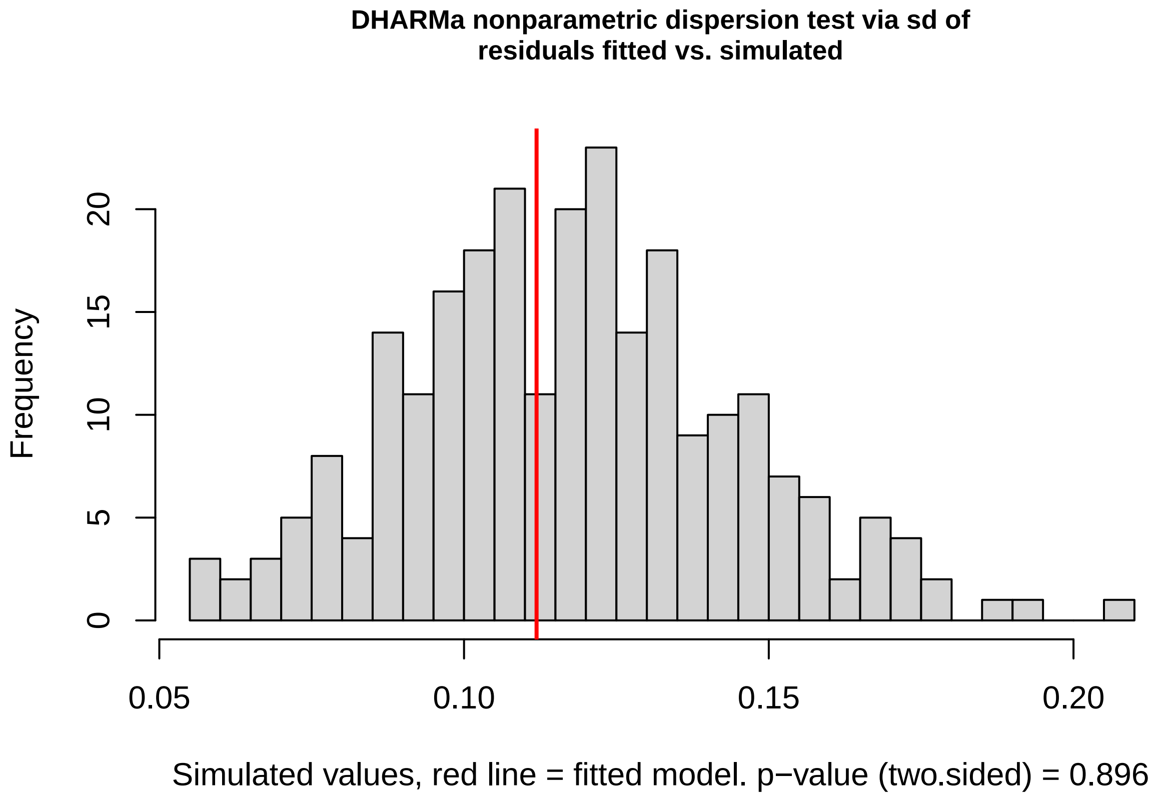

#### 2.2 Power analyses

```
simulate_power <- function(model,
                           effects_pct = 1:20,
                           nsim = 5000,
                           alpha = 0.05) {

  X <- model.matrix(model)
  beta_original <- coef(model)
  sigma_resid <- sigma(model)
  df <- df.residual(model)

  treatment_index <- which(names(beta_original) == "treatVIE")

  if (length(treatment_index) != 1) {
    stop("Could not identify the treatVIE coefficient.")
  }

  # Set the treatment effect to zero
  beta_null <- beta_original
  beta_null[treatment_index] <- 0

  mu_null <- as.vector(X %*% beta_null)

  # Mean predicted CONTROL outcome
  control_mean <- mean(mu_null)

  # Check the scale and value used for the percentage calculation
```

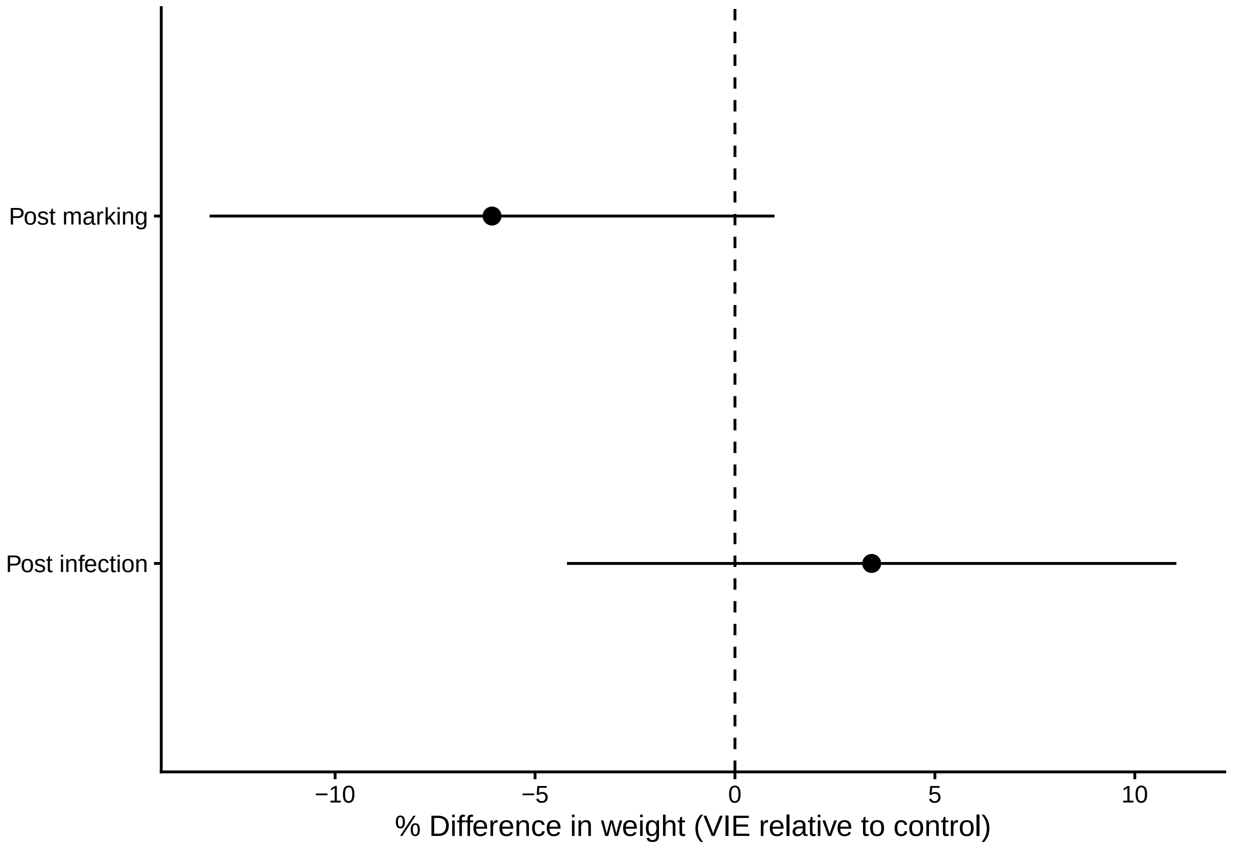

Figure S1: Change in weight did not significantly differ between VIE marked and control fish in either the post injection or post infection time periods. Points give the estimated marginal means from the models, and error bars the 95% confidence intervals.

```r
  print(control_mean)

  results <- data.frame(
    effect_pct = effects_pct,
    power = NA_real_
  )

  vie <- X[, treatment_index]

  for (i in seq_along(effects_pct)) {

    # Convert percentage to an absolute difference
    effect_absolute <- abs(control_mean) * effects_pct[i] / 100

    p_values <- replicate(nsim, {

      y_sim <- mu_null + vie * effect_absolute +
        rnorm(nrow(X), mean = 0, sd = sigma_resid)

      fit_sim <- lm.fit(X, y_sim)

      residual_variance <- sum(fit_sim$residuals^2) / df
      vcov_sim <- residual_variance * solve(crossprod(X))

      estimate <- fit_sim$coefficients[treatment_index]
      se <- sqrt(vcov_sim[treatment_index, treatment_index])

      2 * pt(
        abs(estimate / se),
        df = df,
        lower.tail = FALSE
      )
    })

    results$power[i] <- mean(p_values < alpha)
  }

  results
}
####################

set.seed(123)

power_weight_pre <- simulate_power(m_weight_preinfection)
```

```
## [1] 0.1235701
```

```r
power_length_pre <- simulate_power(m_length_preinfection)
```

```
## [1] 18.10913
```

```r
power_weight_infection <- simulate_power(m_weight_infection)
```

```
## [1] 0.1295911
```

```r
power_length_infection <- simulate_power(m_length_infection)
```

```
## [1] 18.97643
```

```r
###################

power_weight_pre$outcome <- "Weight"
power_weight_pre$period <- "Post marking"

power_length_pre$outcome <- "Length"
power_length_pre$period <- "Post marking"

power_weight_infection$outcome <- "Weight"
power_weight_infection$period <- "Post infection"

power_length_infection$outcome <- "Length"
power_length_infection$period <- "Post infection"

power_results <- rbind(
  power_weight_pre,
  power_length_pre,
  power_weight_infection,
  power_length_infection
)

power_results <- power_results[
  , c("period", "outcome", "effect_pct", "power")
]

####################

library(dplyr)

mde_results <- power_results %>%
  group_by(period, outcome) %>%
  summarise(
    MDE_80pct = if (any(power >= 0.80)) {
      min(effect_pct[power >= 0.80])
    } else {
      NA_real_
    },
    .groups = "drop"
  )

mde_results
```

```
## # A tibble: 4 x 3
##   period         outcome MDE_80pct
##   <chr>          <chr>       <int>
## 1 Post infection Length          5
## 2 Post infection Weight         11
## 3 Post marking   Length          5
## 4 Post marking   Weight         11
```

```
## Warning in geom_vline(data = thresholds, aes(xintercept = effect_pct), colour =
```

```
## "red", : Ignoring unknown parameters: `inherit.aes`
```

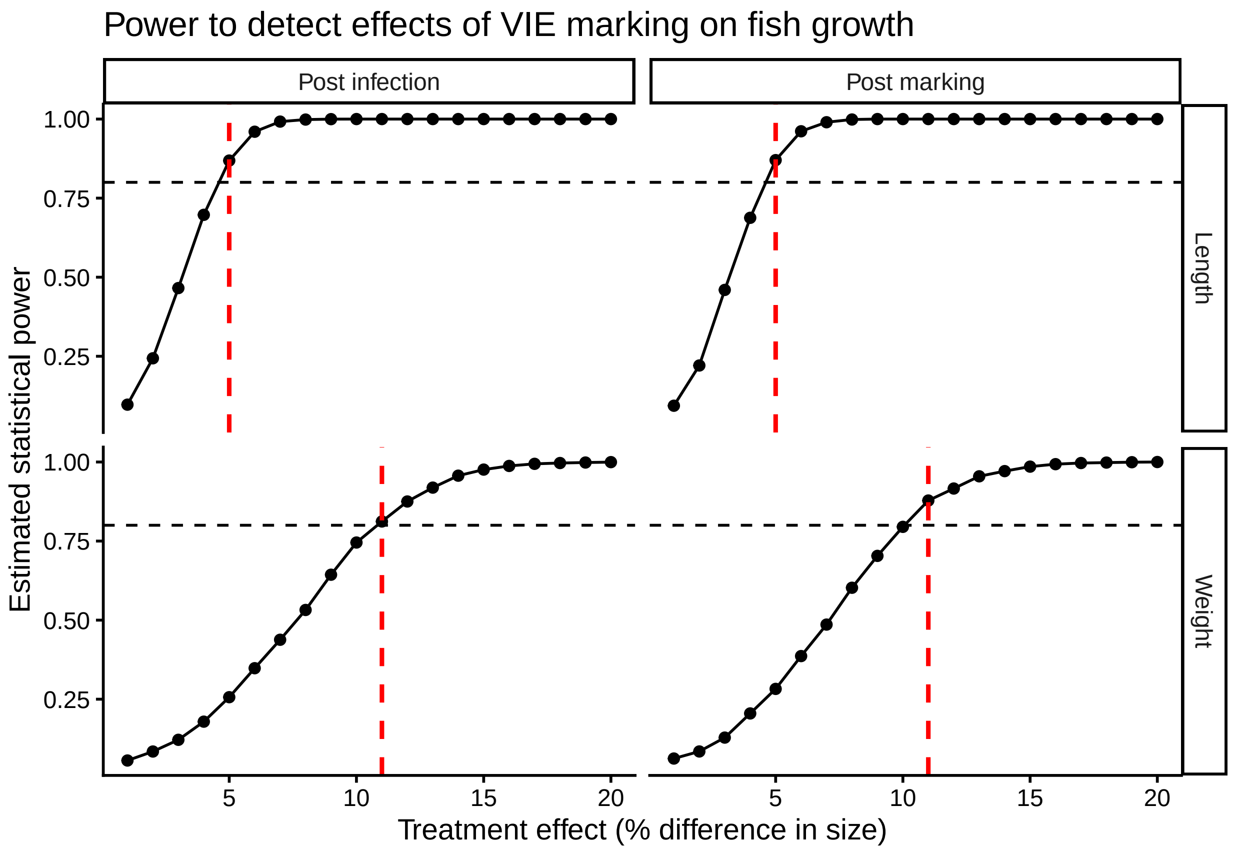

Figure S2: Power analysis shows that the power to detect differences in length exceeded 80% for differences of around 5% across both time periods. However, I had less than 80% power to detect differences of below ~13% in weight across both periods.

### 3 Do worms transmit differently to fish in different treatment groups?

#### 3.1 Model

This model shows that worms transmitted significantly faster to VIE-marked fish than to the controls.

```
minf<-glmer(timeinfs~treat+sex+donorsex+(1|donorid),
        family=gaussian(link="log"),
        data=fdf)

summary(minf)
```

```
## Generalized linear mixed model fit by maximum likelihood (Laplace
##   Approximation) [glmerMod]
##  Family: gaussian  ( log )
## Formula: timeinfs ~ treat + sex + donorsex + (1 | donorid)
##    Data: fdf
##
##      AIC      BIC   logLik deviance df.resid
```

```
##    783.3    794.5   -385.7    771.3       42
##
## Scaled residuals:
##     Min      1Q  Median      3Q     Max
## -2.3982 -0.3868 -0.0646  0.3344  4.4263
##
## Random effects:
##  Groups   Name        Variance Std.Dev.
##  donorid  (Intercept)  7005     83.7
##  Residual             22194    149.0
## Number of obs: 48, groups:  donorid, 13
##
## Fixed effects:
##             Estimate Std. Error t value Pr(>|z|)
## (Intercept)  5.71982    0.25961  22.033   <2e-16 ***
## treatVIE    -0.59704    0.28256  -2.113   0.0346 *
## sexM        -0.07142    0.22270  -0.321   0.7484
## donorsexM   -0.22361    0.31674  -0.706   0.4802
## ---
## Signif. codes:  0 '***' 0.001 '**' 0.01 '*' 0.05 '.' 0.1 ' ' 1
##
## Correlation of Fixed Effects:
##           (Intr) trtVIE sexM
## treatVIE  -0.135
## sexM      -0.430 -0.115
## donorsexM -0.665 -0.130  0.110
```

```
Anova(minf)
```

```
## Analysis of Deviance Table (Type II Wald chisquare tests)
##
## Response: timeinfs
##            Chisq Df Pr(>Chisq)
## treat     4.4646  1    0.03461 *
## sex       0.1028  1    0.74845
## donorsex  0.4984  1    0.48022
## ---
## Signif. codes:  0 '***' 0.001 '**' 0.01 '*' 0.05 '.' 0.1 ' ' 1
```

```
simf<-simulateResiduals(minf, plot=T)
```

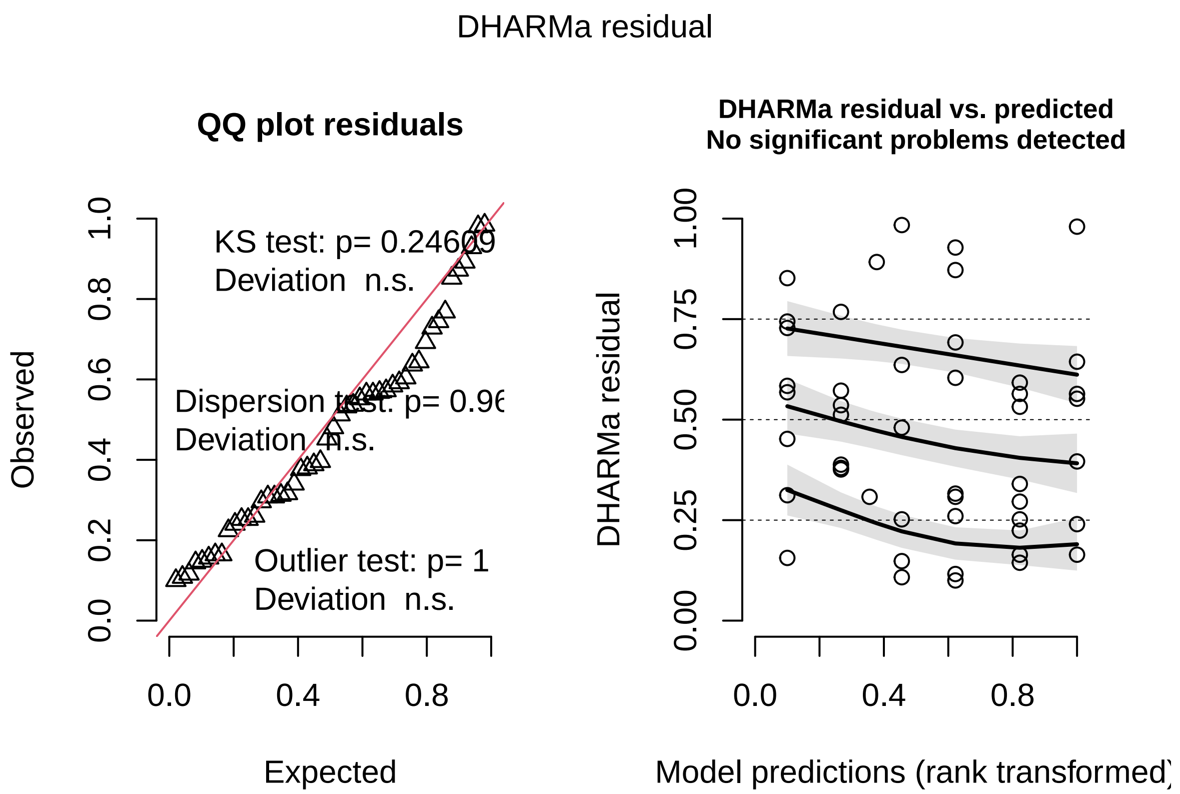

```
testDispersion(simf)
```

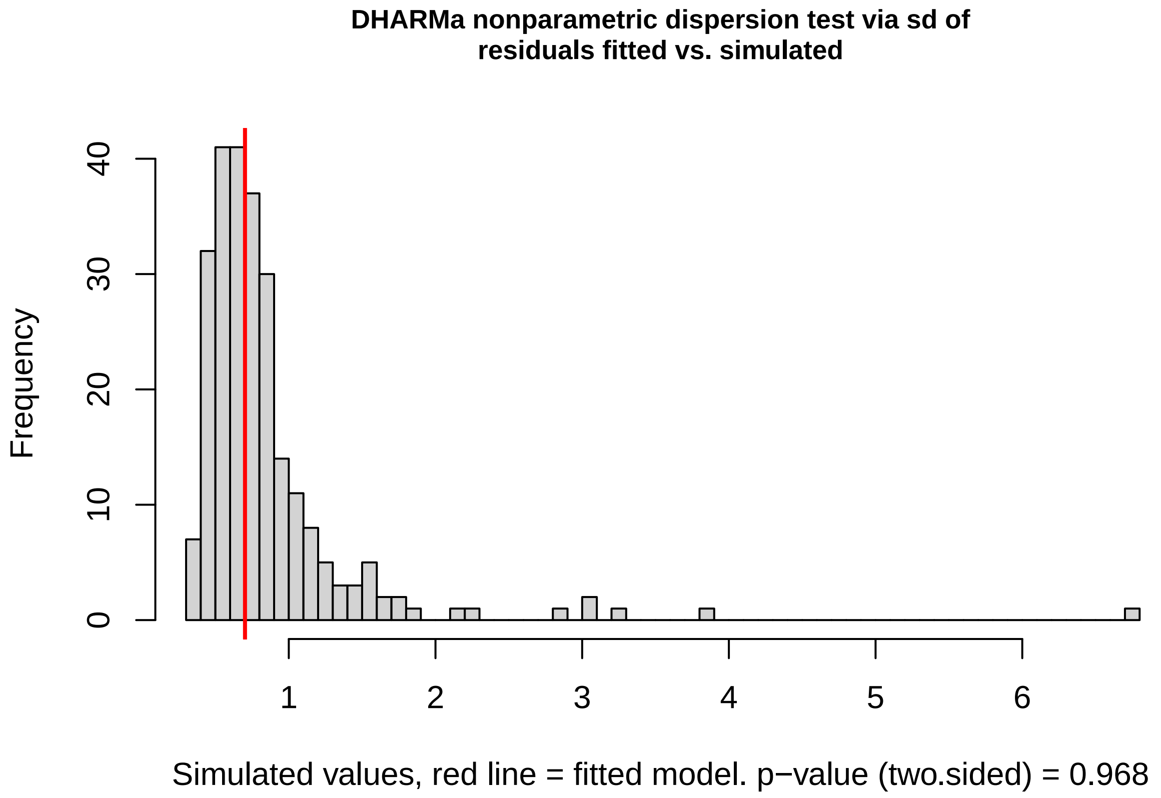

#### 3.2 Effect size

```
emm <- emmeans(minf, ~ treat, type="response")
pairs(emm, reverse = TRUE)
```

```
##  contrast      ratio    SE  df null z.ratio p.value
##  VIE / CONTROL  0.55 0.156 Inf    1  -2.113  0.0346
##
## Results are averaged over the levels of: sex, donorsex
## Tests are performed on the log scale
```

#### 3.3 Leave one out analysis

The result of the model is robust to the removal of donors. The design of the experiment - that each donor contributed worms to multiple experimental fish, haphazard with respect to marking treatment - means that the removal of some donors affects the sample size disproportionately. The removal of four of the donors results in the confidence interval of the effect of marking treatment including 1 - i.e., no significant difference between marking groups. The removal of the remaining 9 donors, however, does not change the significance of the result presented in the main text.

```
library(lme4)
library(dplyr)
library(broom.mixed)
```

```
## Warning: package 'broom.mixed' was built under R version 4.4.3
```

```
dat <- fdf
donors <- unique(dat$donorid)

loo_latency <- list()

for (i in seq_along(donors)) {

  dat_loo <- droplevels(
    dat[dat$donorid != donors[i], ]
  )

  fit <- tryCatch(
    glmer(
      timeinfs ~ treat + sex + donorsex + (1 | donorid),
      family = gaussian(link = "log"),
      data = dat_loo
    ),
    error = function(e) NULL
  )

  if (is.null(fit)) next

  coefs <- summary(fit)$coefficients

  if (!"treatVIE" %in% rownames(coefs)) next

  estimate <- coefs["treatVIE", "Estimate"]
  se <- coefs["treatVIE", "Std. Error"]
```

```r
  loo_latency[[i]] <- data.frame(
    omitted_donor = as.character(donors[i]),
    estimate = estimate,
    ratio = exp(estimate),
    lower_CI = exp(estimate - 1.96 * se),
    upper_CI = exp(estimate + 1.96 * se)
  )
}

loo_latency <- bind_rows(loo_latency)

loo_latency
```

```
##    omitted_donor   estimate     ratio  lower_CI  upper_CI
## 1    W_L3A_5 (M) -0.2756374 0.7590882 0.5209028 1.1061848
## 2    W_L3E_1 (M) -0.5240483 0.5921186 0.3418970 1.0254683
## 3    W_L3B_2 (M) -0.8220901 0.4395121 0.2435512 0.7931426
## 4    W_L3A_9 (F) -0.6414207 0.5265438 0.2859518 0.9695635
## 5    W_L3A_2 (F) -0.5760325 0.5621242 0.3306790 0.9555600
## 6    F_L3C_1 (M) -0.7012430 0.4959684 0.2654659 0.9266148
## 7    F_L3A_1 (M) -0.5841255 0.5575933 0.3082408 1.0086602
## 8    F_L3D_1 (M) -0.6670671 0.5132116 0.2728278 0.9653933
## 9        F_L3C_2 -0.7445236 0.4749605 0.2583412 0.8732153
## 10   F_L3B_1 (M) -0.5977808 0.5500309 0.3000397 1.0083134
## 11   F_L3C_3 (F) -0.6196002 0.5381596 0.3020147 0.9589457
## 12  F_L3OG_2 (F) -0.6041740 0.5465256 0.3065820 0.9742590
## 13  F_L3OG_1 (F) -0.4035955 0.6679142 0.3890570 1.1466429
```

```r
loo_latency <- loo_latency %>%
  mutate(
    ratio_vie_control = ratio,
    lower_vie_control = lower_CI,
    upper_vie_control = upper_CI
  )

loo_summary_latency <- loo_latency %>%
  summarise(
    n = n(),
    min_ratio = min(ratio_vie_control, na.rm = TRUE),
    median_ratio = median(ratio_vie_control, na.rm = TRUE),
    max_ratio = max(ratio_vie_control, na.rm = TRUE),
    min_lower_CI = min(lower_vie_control, na.rm = TRUE),
    max_lower_CI = max(lower_vie_control, na.rm = TRUE),
    min_upper_CI = min(upper_vie_control, na.rm = TRUE),
    max_upper_CI = max(upper_vie_control, na.rm = TRUE),
    .groups = "drop"
  )

loo_summary_latency
```

```
##    n min_ratio median_ratio max_ratio min_lower_CI max_lower_CI min_upper_CI
## 1 13 0.4395121    0.5465256 0.7590882    0.2435512    0.5209028    0.7931426
##   max_upper_CI
## 1     1.146643
```

### 4 Do worms grow differently on fish in different treatment groups?

#### 4.1 Create infection integral metric

```
library(dplyr)

# Function to calculate area under the curve
calc_auc <- function(day, totworm) {

  # Remove missing observations and sort by day
  dat <- data.frame(day = day, totworm = totworm) %>%
    filter(!is.na(day), !is.na(totworm)) %>%
    arrange(day)

  # Need at least two observations
  if (nrow(dat) < 2) return(NA_real_)

  # Trapezoidal rule
  sum(diff(dat$day) *
        (head(dat$totworm, -1) +
           tail(dat$totworm, -1)) / 2)
}

# Calculate AUC for each fish
auc_data <- df0 %>%
  group_by(fishID, treat, lag, sex) %>%
  summarise(
    AUC = calc_auc(day, totworm),
    n_obs = sum(!is.na(day) & !is.na(totworm)),
    first_day = min(day[!is.na(day) & !is.na(totworm)]),
    last_day = max(day[!is.na(day) & !is.na(totworm)]),
    dur = last_day-first_day,
    .groups = "drop"
  )
```

#### 4.2 Model

```
mauc<-glm(AUC~lag*treat+sex+dur, data=auc_data, family=Gamma(link="log"))
summary(mauc)
```

```
##
## Call:
## glm(formula = AUC ~ lag * treat + sex + dur, family = Gamma(link = "log"),
##     data = auc_data)
##
## Coefficients:
##                 Estimate Std. Error t value Pr(>|t|)
## (Intercept)      2.45335    0.72455   3.386  0.00123 **
## lag2             2.86570    0.35034   8.180 1.73e-11 ***
## treatVIE         1.30217    0.48027   2.711  0.00863 **
## sexM             0.33638    0.22773   1.477  0.14463
## dur              0.08850    0.03836   2.307  0.02434 *
## lag2:treatVIE   -1.51033    0.54471  -2.773  0.00730 **
## ---
```

```
## Signif. codes:  0 '***' 0.001 '**' 0.01 '*' 0.05 '.' 0.1 ' ' 1
##
## (Dispersion parameter for Gamma family taken to be 0.8560547)
##
##     Null deviance: 160.51  on 68  degrees of freedom
## Residual deviance: 111.31  on 63  degrees of freedom
## AIC: 1025.5
##
## Number of Fisher Scoring iterations: 18
```

```
Anova(mauc)
```

```
## Analysis of Deviance Table (Type II tests)
##
## Response: AUC
##           LR Chisq Df Pr(>Chisq)
## lag         34.874  1  3.518e-09 ***
## treat        0.076  1    0.78322
## sex          2.130  1    0.14442
## dur          1.785  1    0.18150
## lag:treat    6.277  1    0.01223 *
## ---
## Signif. codes:  0 '***' 0.001 '**' 0.01 '*' 0.05 '.' 0.1 ' ' 1
```

```
sim <- simulateResiduals(mauc)
plot(sim)
```

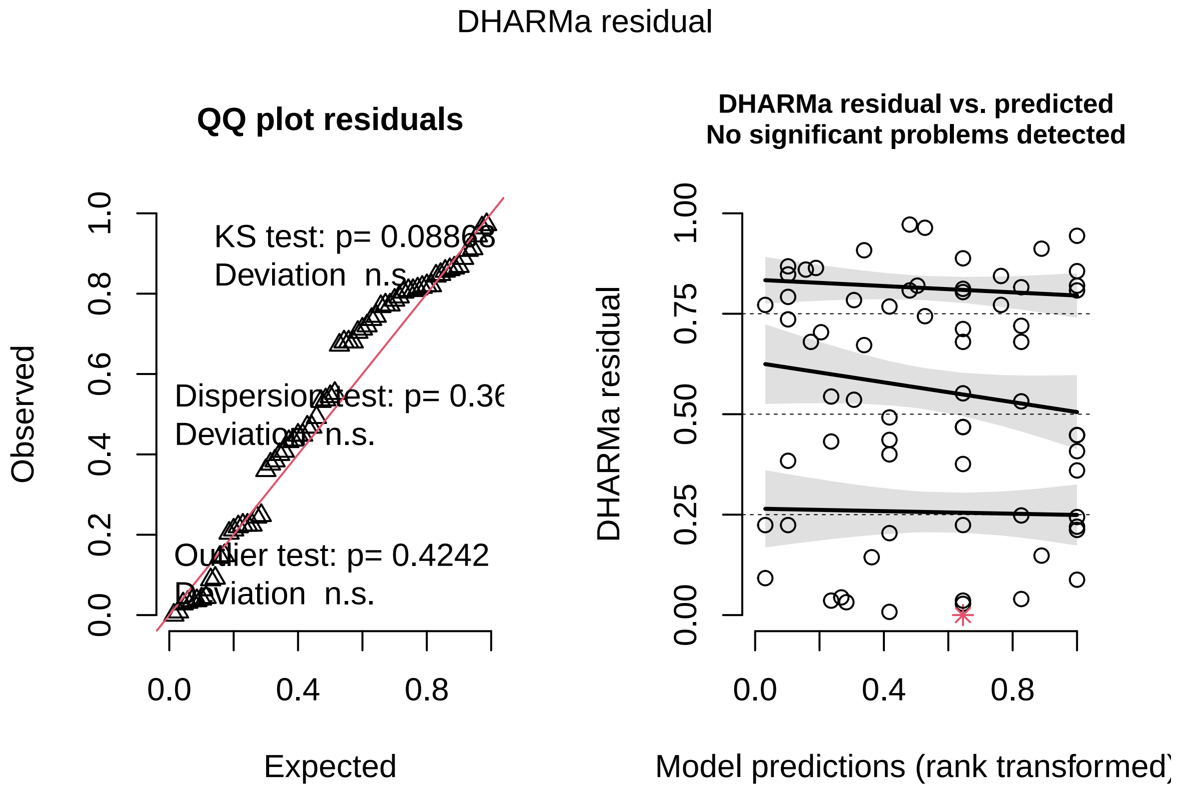

```
testDispersion(sim)
```

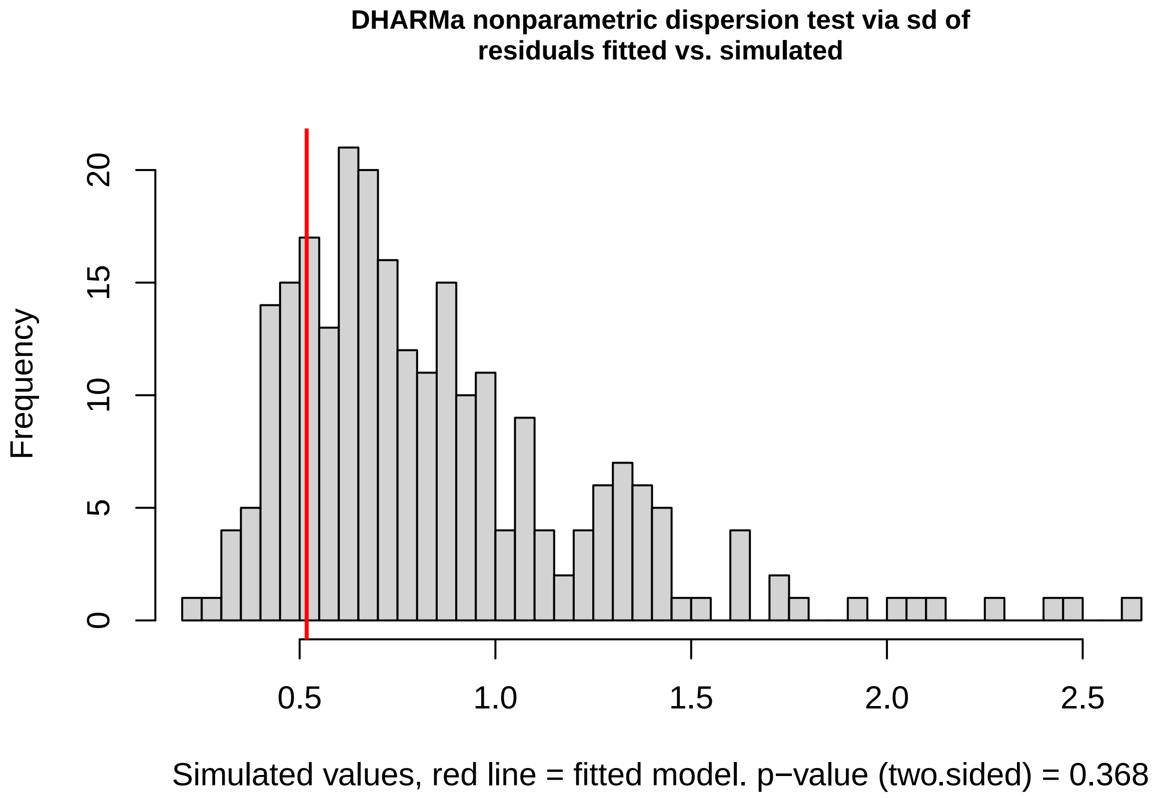

#### 4.3 Effect size

```
emm <- emmeans(mauc, ~ treat | lag)
pairs(emm, reverse = TRUE, type = "response")
```

```
## lag = 1:
##  contrast       ratio    SE df null t.ratio p.value
##  VIE / CONTROL 3.677 1.766 63    1   2.711  0.0086
##
## lag = 2:
##  contrast       ratio    SE df null t.ratio p.value
##  VIE / CONTROL 0.812 0.216 63    1  -0.783  0.4364
##
## Results are averaged over the levels of: sex
## Tests are performed on the log scale
```

#### 4.4 Leave one out analysis

Here, I conducted a 'leave one out' analysis to test how sensitive the results were to the presence of any one fish. The estimated treatment effect at one month post-marking was robust to the exclusion of individual fish, with VIE-to-control AUC ratios ranging from 2.63 to 4.92 across leave-one-fish-out analyses. The lower confidence limit was always above 1: VIE marking was associated with higher infection intensities across leave-one-fish-out analyses. At two months post-marking, ratios ranged from 0.68 to 0.88, and the confidence interval always included 1: the effect of VIE marking on infection intensity was not significant in any of the leave-one-fish-out analyses. Thus, the direction of the estimated treatment effect across lag groups was unchanged by the exclusion of any single fish.

```
#AUC dataset
dat <- auc_data
```

```r
fish_ids <- unique(dat$fishID)

loo_results_auc <- list()

for (i in seq_along(fish_ids)) {

  message("Leaving out fish ", fish_ids[i],
          " (", i, " of ", length(fish_ids), ")")

  # Remove all data for one fish
  dat_loo <- droplevels(
    dat[dat$fishID != fish_ids[i], ]
  )

  # Refit the original AUC model
  fit_loo <- tryCatch(
    glm(
      AUC ~ lag * treat + sex + dur,
      data = dat_loo,
      family = family(mauc)
    ),
    error = function(e) {
      message("Model failed: ", conditionMessage(e))
      NULL
    }
  )

  if (is.null(fit_loo)) next

  # Treatment comparisons within each lag
  result <- tryCatch({

    emm <- emmeans(fit_loo, ~ treat | lag)

    contrasts <- contrast(
      emm,
      method = "revpairwise"
    )

    res <- as.data.frame(
      summary(
        contrasts,
        type = "response",
        infer = c(TRUE, TRUE)
      )
    )

    res$omitted_fish <- as.character(fish_ids[i])
    res

  }, error = function(e) {
    message("Contrast failed: ", conditionMessage(e))
    NULL
```

```r
  })

  if (!is.null(result)) {
    loo_results_auc[[length(loo_results_auc) + 1]] <- result
  }
}

# Combine successful fits
loo_results_auc <- bind_rows(loo_results_auc)

loo_results_auc <- loo_results_auc %>%
  mutate(
    ratio_vie_control = ratio,
    lower_vie_control = lower.CL,
    upper_vie_control = upper.CL
  )

loo_summary_auc <- loo_results_auc %>%
  group_by(lag) %>%
  summarise(
    n = n(),
    min_ratio = min(ratio_vie_control, na.rm = TRUE),
    median_ratio = median(ratio_vie_control, na.rm = TRUE),
    max_ratio = max(ratio_vie_control, na.rm = TRUE),
    min_lower_CI = min(lower_vie_control, na.rm = TRUE),
    max_lower_CI = max(lower_vie_control, na.rm = TRUE),
    min_upper_CI = min(upper_vie_control, na.rm = TRUE),
    max_upper_CI = max(upper_vie_control, na.rm = TRUE),
    .groups = "drop"
  )

loo_summary_auc
```

```
## # A tibble: 2 x 9
##   lag       n min_ratio median_ratio max_ratio min_lower_CI max_lower_CI
##   <fct> <int>     <dbl>        <dbl>     <dbl>        <dbl>        <dbl>
## 1 1        69     2.63         3.69      4.92         1.02         1.78
## 2 2        69     0.682        0.813     0.882        0.406        0.519
## # i 2 more variables: min_upper_CI <dbl>, max_upper_CI <dbl>
```

### 5 R and Library Versions

```r
version
```

```
##                _
## platform       aarch64-apple-darwin20
## arch           aarch64
## os             darwin20
## system         aarch64, darwin20
## status
## major          4
## minor          4.0
```

```
## year           2024
## month          04
## day            24
## svn rev        86474
## language       R
## version.string R version 4.4.0 (2024-04-24)
## nickname       Puppy Cup
```

```r
installed.packages()[names(sessionInfo()$otherPkgs), "Version"]
```

```
## broom.mixed        mgcv        nlme     lmodel2        lme4      Matrix
##   "0.2.9.7"     "1.9-4"   "3.1-164"     "1.7-4"  "1.1-35.3"     "1.7-0"
##        MASS     emmeans   patchwork  kableExtra       knitr   lubridate
##    "7.3-64"    "1.10.1"     "1.3.2"     "1.4.0"      "1.51"     "1.9.4"
##     forcats     stringr       dplyr       purrr       readr       tidyr
##     "1.0.0"     "1.5.1"     "1.2.0"     "1.2.1"     "2.1.5"     "1.3.2"
##      tibble   tidyverse      visreg      DHARMa         car     carData
##     "3.2.1"     "2.0.0"     "2.7.0"     "0.4.7"     "3.1-2"     "3.0-5"
##     glmmTMB      GGally     ggplot2      readxl        here
##     "1.1.9"     "2.4.0"     "4.0.0"     "1.4.3"     "1.0.1"
```
